# Interactions with Commensal and Pathogenic Bacteria Induce HIV-1 Latency in Macrophages through Altered Transcription Factor Recruitment to the LTR

**DOI:** 10.1101/2020.05.18.103333

**Authors:** Gregory A. Viglianti, Vicente Planelles, Timothy M. Hanley

## Abstract

Macrophages are infected by HIV-1 *in vivo* and contribute to both viral spread and pathogenesis. Recent human and animal studies suggest that HIV-1-infected macrophages serve as a reservoir that contributes to HIV-1 persistence during anti-retroviral therapy. The ability of macrophages to serve as persistent viral reservoirs is likely influenced by the local tissue microenvironment, including interactions with pathogenic and commensal microbes. Here we show that the sexually transmitted pathogen *Neisseria gonorrhoeae* (GC) and the gut-associated microbe *Escherichia coli (E. coli),* which encode ligands for both Toll-like receptor 2 (TLR2) and TLR4, repressed HIV-1 replication in macrophages and thereby induced a state reminiscent of viral latency. This repression was mediated by signaling through TLR4 and the adaptor protein TRIF and was associated with increased production of type I interferons. Inhibiting TLR4 signaling, blocking type 1 interferon, or knocking-down TRIF reversed LPS- and GC-mediated repression of HIV-1. Finally, the repression of HIV-1 in macrophages was associated with the recruitment of interferon regulatory factor 8 (IRF8) to the interferon stimulated response element (ISRE) downstream of the 5′ HIV-1 long terminal repeat (LTR). Our data indicate that IRF8 is responsible for repression of HIV-1 replication in macrophages in response to TRIF-dependent signaling during GC and *E. coli* co-infection. These findings highlight the potential role of macrophages as HIV-1 reservoirs as well as the role of the tissue microenvironment and co-infections as modulators of HIV-1 persistence.

**IMPORTANCE:** The major barrier toward the eradication of HIV-1 infection is the presence of a small reservoir of latently infected cells, which include CD4+ T cells and macrophages that escape immune-mediated clearance and the effects of anti-retroviral therapy. There remain crucial gaps in our understanding of the molecular mechanisms that lead to transcriptionally silent or latent HIV-1 infection of macrophages. The significance of our research is in identifying microenvironmental factors, such as commensal and pathogenic microbes, that can contribute to the establishment and maintenance of latent HIV-1 infection in macrophages. It is hoped that identifying key processes contributing to HIV-1 persistence in macrophages may ultimately lead to novel therapeutics to eliminate latent HIV-1 reservoirs *in vivo.*

## INTRODUCTION

Macrophages are among the immune cells located within the gastrointestinal and genitourinary mucosae thought to play a role in HIV-1 sexual transmission and pathogenesis (1–3). A number of studies examining either HIV-1 infection of human vaginocervical or gastrointestinal tissue explants or SIV_mac_ infection in rhesus macaque animal models have shown that macrophages are among the earliest cells infected during mucosal transmission (2, 4, 5). Macrophages can be productively infected with HIV-1 and are thought to be a source of virus persistence *in vivo* (6). Given their role in transmission, pathogenesis, and viral persistence, it is important to understand how the local mucosal microenvironment and cellular signaling pathways modulate interactions between macrophages and HIV-1.

Sexually-transmitted infections (STIs) have been shown to be co-factors that enhance HIV-1 transmission (7). *Neisseria gonorrhoeae* (gonococcus, GC) is a non-ulcerative STI that is thought to augment mucosal transmission of HIV-1 both by inducing inflammation and by directly activating virus infection and replication (8–13). The role of GC in HIV-1 persistence is less well understood. Several studies have implicated GC-encoded pathogen-associated molecular patterns (PAMPs) as mediators of both inflammation and HIV-1 activation in target cells such as macrophages; however, the interactions between GC and macrophages are complex. GC encodes PAMPs capable of engaging Toll-like receptors (TLRs), including TLR2, TLR4, and TLR9 (14, 15). While the effects of co-infection with live GC on HIV-1 replication in macrophages have not been reported, purified lipooligosaccharide (LOS) as well as *Escherichia coli* lipopolysaccharide (LPS) have been shown to repress virus replication through the production of type 1 interferons (IFNs) (16, 17). In the case of LPS, repression is due to undefined effects at the level of gene expression. Although it is not entirely clear how TLR2 signaling affects HIV-1 expression in macrophages, studies have shown that purified TLR2 ligands activate virus replication in macrophages (18) and latently-infected T cells (19).

Here, we demonstrate that co-infection with GC and *E. coli* repress HIV-1 expression in macrophages. To investigate the underlying mechanism(s) responsible for this repression, we examined the individual effects of TLR2 and TLR4 signaling on HIV-1 expression in macrophages. TLR2 signaling activated HIV-1 expression in macrophages, whereas TLR4 signaling repressed virus expression. Importantly, TLR4 signaling overcame the activation effects of TLR2 signaling in macrophages. The TLR4-mediated repression of HIV-1 in macrophages co-infected with GC or *E. coli* was dependent on signaling through Toll/IL-1 receptor domain-containing adapter inducing interferon-β (TRIF) and required type 1 IFN production. Finally, we showed that TLR4 signaling leads to the late-phase recruitment of IRF8 to the interferon-stimulated response element (ISRE) downstream of the 5′ HIV-1 LTR in infected macrophages. Taken together, our data suggest TRIF-mediated signaling represses HIV-1 replication in response to GC or *E. coli* co-infection in an IRF8-dependent manner and shifts macrophages from a state of robust HIV-1 expression to a state of persistent low-level/latent infection.

## MATERIALS AND METHODS

### Ethics Statement

This research has been determined to be exempt by the Institutional Review Board of the Boston University Medical Center since it does not meet the definition of human subjects research.

### Cell isolation and culture

Primary human CD14+ monocytes were isolated from the peripheral blood mononuclear cells of healthy donors using anti-CD14 magnetic beads (Miltenyi Biotec) per the manufacturer’s instructions. Primary monocyte-derived macrophages (MDMs) were generated by culturing CD14+ monocytes in the presence of 10% human AB serum and 10% FBS for 6 days. Following differentiation, MDMs were cultured in RPMI-1640 supplemented with 10% FBS, 100 U/ml penicillin, 100 μg/ml streptomycin, and 0.29 mg/ml L-glutamine. The genetic sex of a subset of the donors was determined by PCR amplification of the SRY gene located on the Y chromosome. PM1 cells were cultured in RPMI-1640 supplemented with 10% FBS, 100 U/ml penicillin, 100 μg/ml streptomycin, and 0.29 mg/ml L-glutamine. 293T cells were cultured in DMEM supplemented with 10% FBS, 100 U/ml penicillin, 100 μg/ml streptomycin, and 0.29 mg/ml L-glutamine. MAGI-CCR5 cells were cultured in DMEM supplemented with 10% FBS, 100U/ml penicillin, 100 μg/ml streptomycin, 0.29 mg/ml L-glutamine, 500 μg/ml G418, 1 μg/ml puromycin, and 0.1 μg/ml hygromycin B. HEK293-TLR2^CFP^TLR1^YFP^ cells and HEK293-TLR4^CFP^/MD-2/CD14 cells were cultured in DMEM supplemented with 10% FBS, 10 μg/ml ciprofloxacin, 0.29 mg/ml L-glutamine, and 500 μg/ml G418.

### Bacterial culture

*Neisseria gonorrhea* (GC) strain FA1090B was a generous gift from Dr. Caroline Genco. GC was cultured from a glycerol stock on GC agar plates supplemented with IsoVitalex Enrichment (Becton Dickinson) in a humidified 37°C incubator with 5% CO_2_. *E. coli* strain DH5α was purchased from New England Biolabs and was cultured from a glycerol stock on LB agar plates at 37°C. Where indicated, bacteria were heat inactivated (heat killed) by incubation at 56°C for 2 hours. Heat inactivation was monitored by culture on GC or LB agar plates as described above.

### Flow cytometry

TLR expression on viable MDMs was assessed eight days after isolation using antibodies against TLR2 (clone TL2.1) and TLR4 (clone HTA125) (both from eBioscience) and eFluor 450 fixable viability dye (eBioscience). MDMs were stained in plates, washed with phosphate-buffered saline (PBS), fixed using BD Cytofix (BD Biosciences), and then detached after incubation in PBS supplemented with 20 mM EDTA for 1 hour at 4°C. Flow cytometric data was acquired using a Becton-Dickenson FACScan II or LSRFortessa and data was analyzed using FlowJo software.

### TLR ligands, interferons, and chemical inhibitors

PAM3CSK4, FSL-1, *Salmonella typhimurium* flagellin (FLA-ST), poly I:C, and *E*. *coli* K12 LPS were obtained from Invivogen. TLR ligands were reconstituted in endotoxin-free H_2_O. IFN-α and IFN-β were purchased from PBL Interferon Source. B18R was purchased from Abcam. BAY 11-7082, celastrol, U0126, PD95809, and SB203580 were purchased from Sigma and reconstituted in DMSO. Dynasore was purchased from Tocris Bioscience and was reconstituted in DMSO.

### Virus Production

Single-round replication-defective HIV-1 reporter viruses were generated by packaging a luciferase expressing reporter virus, BruΔEnvLuc2, or an enhanced green fluorescent protein expressing reporter virus, BruΔEnvEGFP3, with the envelope glycoproteins from VSV (VSV-G). In these constructs, reporter gene expression is under the control of the 5′ LTR. Reporter virus stocks were generated by transfecting HEK293T cells using the calcium phosphate method as described previously (18). Replication competent HIV-1_Ba-L_ was generated by infection of PM1 cells as described previously (18). Virus titers were determined using MAGI-CCR5 cells, and p24^gag^ content was determined by ELISA as described previously (18).

### Virus infections

To assess viral replication, macrophages (2.5 × 10^5^ cells/well in 24-well plates) were incubated with VSV-G-pseudotyped HIV-luciferase reporter virus at a multiplicity of infection (MOI) of 0.1 for 4 hours at 37°C. Cells were washed four to five times with PBS to remove unbound virus, and cultured in growth medium. Following 48 hours of culture, cells were treated with TLR ligands or vehicle, as indicated in the text and figure legends. After 18 hours, the cells were washed twice with PBS and lysed in PBS/0.02% Triton X-100. Luciferase activity was measured using BrightGlo luciferase reagent (Promega) and a MSII luminometer.

### HIV-1 transcription

Total cytoplasmic RNA was isolated from MDMs using the RNeasy Mini kit (Qiagen). RNA (100 ng) was analyzed by reverse transcription-PCR (RT-PCR) using the OneStep RT-PCR kit (Qiagen). RNA was reverse transcribed and amplified in a total volume of 50 μl containing 2.5 mM MgCl2, 400 μM concentrations of each deoxynucleoside triphosphate, 10 U of RNasin RNase inhibitor (Promega), 5 μCi of α-^32^P dATP, and 0.6 μM HIV-1 specific primers. RNA samples were reverse transcribed for 30 minutes at 50°C. After an initial denaturing step at 95°C for 15 minutes, cDNA products were amplified for 25 cycles each consisting of a 30-second denaturing step at 94°C, a 45-second annealing step at 65°C, and a one-minute extension step at 72°C. The amplification concluded with a 10-minute extension step at 72°C. Samples were resolved on 5% nondenaturing polyacrylamide gels, visualized by autoradiography, and quantified in a Molecular Dynamics PhosphorImager SI using ImageQuant software. Alternatively, HIV-1 RNA was analyzed using the QuantiTect SYBR Green RT-PCR kit (Qiagen) in a LightCycler 480 (Roche). The HIV-1 primers were specific for the R and U5 regions of the LTR and amplify both spliced mRNAs and genomic RNA. The HIV-1 primers were sense primer 5′-GGCTAACTAGGGAACCCACTGC-3′ and antisense primer 5′-CTGCTAGAGATTTTCCACACTGAC-3′). a-tubulin primers were: sense primer 5′ CACCCGTCTTCAGGGCTTCTTGGTTT-3′ and antisense primer, 5′CATTTCACCATCTGGTTGGCTGGCTC-3′. RNA standards corresponding to 500, 50, and 5 ng of RNA from PAM3CSK4-activated MDMs were included in each experiment to ensure that all amplifications were within the linear range of the assay.

### HIV-1 RNA stability assays

MDMs (2 × 10^6^ cells/well in 6-well plates) were incubated with VSV-G-pseudotyped HIV-luciferase reporter virus at an MOI of 0.1 for 4 h at 37°C. Cells were washed four to five times with PBS to remove unbound virus and cultured in growth medium. Following 48 h of culture, cells were treated with TLR ligands (PAM3CSK4 or LPS at 100 ng/ml) or vehicle for 4 hours. Actinomycin D (10 μg/ml) was then added to cells to block *de novo* RNA synthesis, and total cytoplasmic RNA was isolated at given times as described in the figure legends. Viral RNA was analyzed using the QuantiTect SYBR Green RT-PCR kit (Qiagen) in a LightCycler 480 (Roche) with primers specific for the R and U5 regions of the LTR as described above.

### Cytokine release assays

MDMs (2.5 × 10^5^ cells/well) were treated with PAM3CSK4 (100 ng/ml), LPS (100 ng/ml), or GC (MOI of 10) for 24 hours. Cell-free culture supernatants were collected and analyzed for TNF-α (eBioscience) or IFN-β (PBL Interferon Source) release by commercially-available ELISA following the manufacturer’s instructions.

### Chromatin immunoprecipitation assays

1.2 × 10^7^ MDMs were incubated with VSV-G-pseudotyped HIV-EGFP reporter virus at an MOI of 2 for four hours at 37°C. Cells were washed four to five times with PBS to remove unbound virus and cultured in growth medium. Following 48 hours of culture, MDMs were treated with TLR ligands for various times as described in the text. Cells were then fixed in 1% formaldehyde for 10 minutes at room temperature, quenched with 125 mM glycine, and lysed in SDS lysis buffer (1% SDS, 10 mM EDTA, 50 mM Tris pH 8.1, 1 mM PMSF, 1 μg/ml aprotinin, 1 μg/ml pepstatin A). Cellular lysates were sonicated using a cup horn (550 Sonic Dismembrator, Fisher Scientific) at a power setting of 5 with twenty-five 20-second pulses on ice, which fragmented the chromatin to an average length of approximately 1000 bp. Samples were diluted and immunoprecipitated with antibodies against NF-κB p65, IRF1, IRF2, IRF4, IRF8, rabbit IgG, or goat IgG (all from Santa Cruz Biotechnology). Purified DNA samples from both ChIPs and input controls were resuspended in distilled H_2_O and analyzed by semi-quantitative PCR. PCR reactions contained 10 mm Tris-HCl pH 8.3, 50 mm KCl, 1.5 mm MgCl2, 100 pmol of each primer, 200 μm each dATP, dGTP, dCTP, and dTTP, 5μCi α^32^P-dATP, and 2.5 units of Amplitaq Gold (Applied Biosystems) in a 50-μl reaction volume. Following an initial denaturation step at 95°C for 15 minutes, DNAs were amplified for 30 cycles, each consisting of a 30-second denaturing step at 94°C, a 45-second annealing step at 65°C, and a one-minute extension step at 72°C. Samples were electrophoresed on 5% nondenaturing polyacrylamide gels, visualized by autoradiography, and quantified using a Molecular Dynamics PhosphorImager SI using ImageQuant software. Alternatively, purified DNA from ChIPs and input controls were analyzed using the PowerUp SYBR Green Master Mix (Applied Biosystems) in a LightCycler 480 (Roche). Primers used to amplify specifically the HIV-1 5′ LTR and GLS were 5′-TGGAAGGGCTAATTTACTCCC-3′ (sense) and 5′-CATCTCTCTCCTTCTAGCCTC-3′ (antisense). Control amplifications of a serial dilution of purified genomic DNA from latently infected U1 cells were performed with each primer set to ensure that all amplifications were within the linear range of the reaction. To calculate the relative levels of association with the LTR, PhosphorImager data of the PCR products obtained for immunoprecipitated chromatin samples were normalized against the PCR products obtained for input DNA (% Input). Values were normalized across donors and expressed as relative binding.

### LTR mutant construction

The reported plasmid pLTR(Sp1)-luciferase was generated by PCR amplification of pNL4-3 using the sense primer 5′-CGGGGTACCCCGTGGAAGGGCTAATTTGGTCCC-3′ and the antisense primers 5′-CCGCTCGAGCGGCATCTCTCTCCTTCTAGCCTC-3′, digestion with KpnI and XhoI, and ligation into KpnI/XhoI-digested pGL3-Basic (Promega). Mutations to the NF-κB and IRF binding sites in pLTR(Sp1)-luciferase were generated using the QuickChange IIXL site-directed mutagenesis kit (Stratagene). Primers used for site-directed mutagenesis are listed in Table 1. The −158 LTR-luciferase construct was generated by deleting the LTR sequence upstream of position −158 (relative to the start site of transcription) of pNL4-3, which includes the AP-1 binding sites located in the U3 portion of the 5′ LTR, digestion of the resulting fragment with KpnI and XhoI, and ligation into KpnI/XhoI-digested pGL3-Basic (Promega).

**Table 1.**
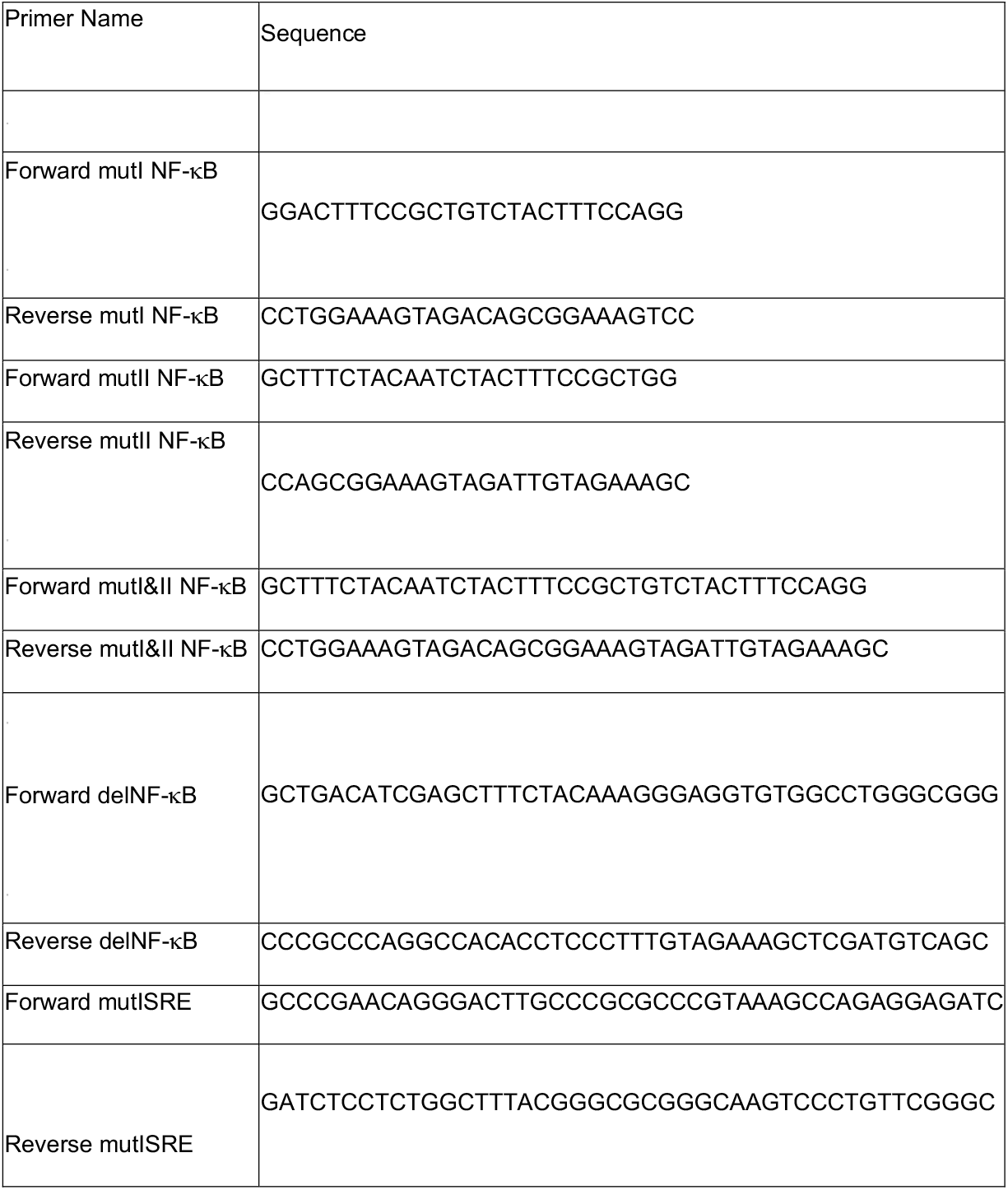
Primers used for PCR-based mutagenesis.

### shRNA knock-down of MyD88, TRIF, and IRF8

MDMs (1.2 × 10^7^) were transfected with plasmids that encoded either a mixture of three to five shRNAs directed against MyD88, a mixture of three to five shRNAs directed against TRIF, or a mixture of three to five control shRNAs (Invivogen) and a blasticidin-resistance gene using Oligofectamine (Invitrogen) per the manufacturer’s instructions. Transfected cells were selected by culture in the presence of blasticidin for 48 hours, and either used in HIV-1 replication assays or lysed for immunoblot analysis to measure MyD88 and TRIF expression using a rabbit monoclonal antibody to MyD88 (Cell Signaling Technology), a rabbit polyclonal antibody to TRIF (Cell Signaling Technology), or a mouse monoclonal antibody to β-actin (Sigma). Similarly, MDMs were transfected with plasmids that encoded either a mixture of three to five shRNAs directed against IRF8 (Sigma) or a mixture of control shRNAs (Sigma) and a puromycin-resistance gene using Oligofectamine (Invitrogen) per the manufacturer’s instructions. Transfected cells were selected by culture in the presence of puromycin for 48 hours and either used in HIV-1 replication assays or lysed for immunoblot analysis to measure IRF8 expression using a rabbit monoclonal antibody (Cell Signaling Technology).

### Overexpression of IRF8

MDMs (1.2 × 10^7^) were transfected with a plasmid that encoded IRF8 (Origene) and a neomycin-resistance gene using Oligofectamine (Invitrogen) per the manufacturer’s instructions. Transfected cells were selected by culture in the presence of neomycin for 48 hours and then used for HIV-1 replication assays or lysed for immunoblot analysis to measure IRF8 expression using a rabbit monoclonal antibody (Cell Signaling Technology).

### Endocytosis/phagocytosis assays

MDMs (5×10^5^/well) were treated with Dynasore (80 μM) or DMSO and then incubated with pHrodo Green *E. coli* particles (Thermo Fisher) at 1 mg/ml for 2 hours at 37°C. The MDMs were then washed three times with PBS, incubated with eFluor 450 fixable viability dye (eBioscience) for 15 minutes at 4°C, and analyzed by flow cytometry. Flow cytometric data was acquired using an Becton-Dickenson LSRFortessa, and data was analyzed using FlowJo software.

### Statistical analysis

Comparison between experimental samples was performed with a paired one-tailed t-test with p < 0.5 denoting significant differences. Experiments were performed in triplicate using cells from a minimum of four independent donors (unless otherwise indicated) to control for interdonor variability.

## RESULTS

### HIV-1 gene expression in MDMs is enhanced or repressed in a TLR-specific manner

To determine how purified TLR ligands affected HIV-1 gene expression, MDMs were infected with a single-round infectious HIV-1 reporter virus, and then treated with ligands for TLR2, TLR3, TLR4, or TLR5. Ligands that activated TLR2 or TLR5 enhanced HIV-1 replication, whereas ligands for TLR3 or TLR4 repressed HIV-1 expression (Figure 1A). The effects of TLR ligands on HIV-1 replication occurred at the level of transcription, as treatment with the TLR2/1 ligand PAM3CSK4 led to an increase in HIV-1 mRNA accumulation, whereas treatment with the TLR4 ligand LPS led to a decrease in HIV-1 transcript levels (Figure 1C-1D). TLR treatment had no effect of viral RNA stability, as viral RNA from LPS-treated MDMs had a similar decay rate to that from untreated MDMs (Figure 1E). Recent studies have demonstrated that myeloid cells from males and females have different susceptibilities to HIV-1 infection, largely due to differential levels of innate immune responses and steroid hormones (20–22). We therefore sought to determine whether there was a sex-based difference in the response to TLR ligand treatment in MDMs. We found that TLR stimulation had similar effects on HIV-1 expression in MDMs from both male and female donors (Supplemental Figure 1). These results indicate that MyD88-dependent signaling enhances HIV-1 transcription whereas TRIF-dependent signaling inhibits HIV-1 transcription in MDMs.

**Figure 1.**
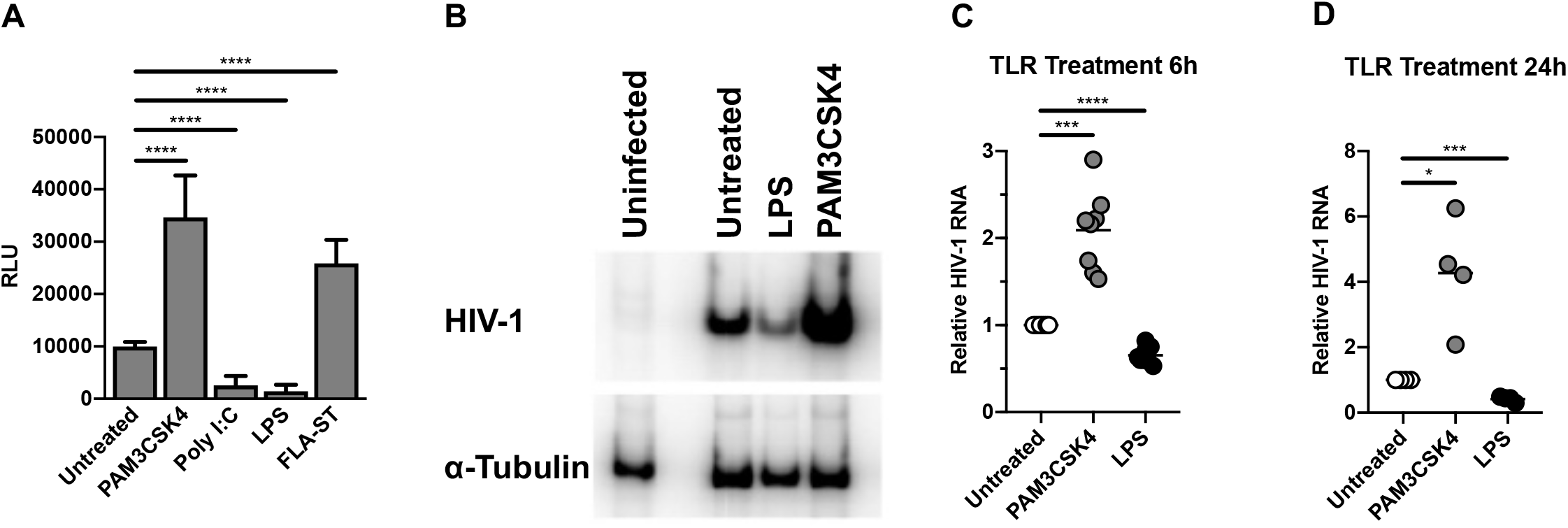
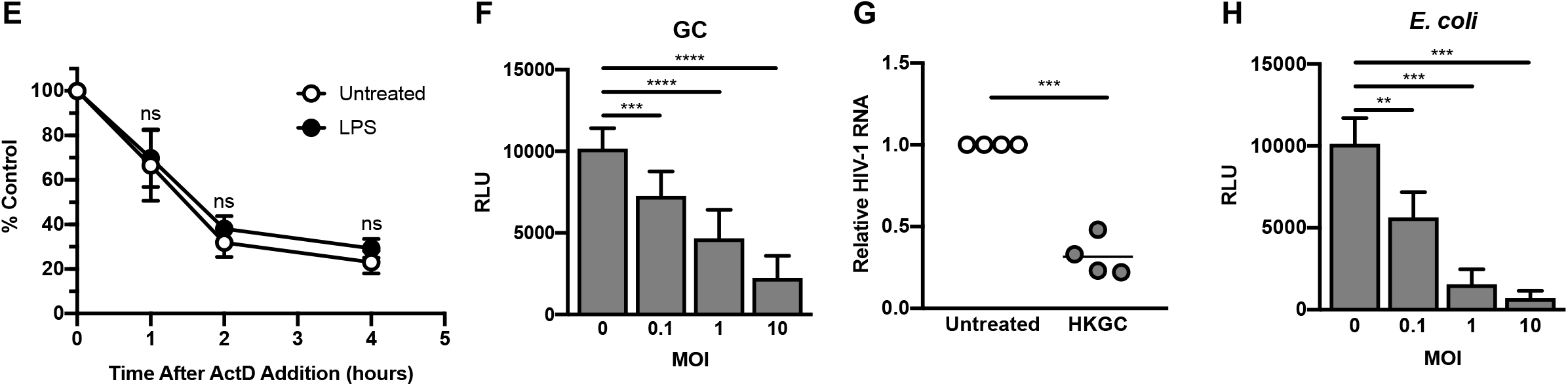
Treatment with purified TLR ligands and intact bacteria alter HIV-1 replication at the level of transcription. (A) MDMs (2.5 × 10^5^ cells/well) were infected with a single-round, replication-defective HIV-luciferase reporter virus at an MOI = 0.1. After four hours, unbound virus was removed by washing with PBS and cells were cultured in complete medium. Fortyeight hours after infection, cells were treated with the TLR2 ligand PAM3CSK4 (100 ng/ml), the TLR3 ligand poly I:C (25 μg/ml), the TLR4 ligand LPS (100 ng/ml), or the TLR5 ligand FLA-ST (100 ng/ml) for 18 hours. The cells were then lysed and assayed for luciferase activity. Bars represent the mean (± SD) of 11 donors, each donor tested in triplicate. (B-D) MDMs were infected as described above. Forty-eight hours after infection, cells were treated with PAM3CSK4 (100 ng/ml) or LPS (100 ng/ml) for six hours (B-C) or 24 hours (D). Cells were then lysed and assayed for viral RNA accumulation by RT-PCR. Shown are data from one representative donor (B) and composite data from eight donors at 6 hours (C) and four donors at 24 hours (D). (E) MDMs (1 × 10^6^ cells/well) were infected as in (A). Forty-eight hours after infection, cells were treated with the TLR4 ligand LPS (100 ng/ml) for 4 h. Cells were then treated with actinomycin D (10 μg/ml) to inhibit transcription. Total cytoplasmic RNA was prepared from the treated cultures at the indicated time points following actinomycin D treatment and analyzed by RT-qPCR for the expression of HIV-1 RNA. The data are the means (± SD) from four donors. (F) MDMs were infected as in (A). Forty-eight hours after infection, the cells were cultured overnight with increasing amounts of GC. Cells were then lysed and assayed for luciferase activity. Bars represent mean (± SD) of seven donors, each donor tested in triplicate. (G) MDMs were infected as described above. Forty-eight hours after infection, cells were treated with GC at an MOI of 10 for 24 hours. Cells were then lysed and assayed for viral RNA accumulation by RT-qPCR. Shown are data from four donors. (H) MDMs (2.5 × 10^5^ cells/well) were infected as in (A). Forty-eight hours after infection, the cells were cultured overnight with increasing amounts of *E. coli.* Cells were then lysed and assayed for luciferase activity. Bars represent mean (± SD) of four donors, each donor tested in triplicate. *, p < 0.05; **, p < 0.01; ***, p < 0.001; p < 0.0001.

MyD88-dependent TLR signaling leads to the activation of both NF-κB and AP-1 transcription factors, among others (23). The HIV-1 LTR contains binding sites for both NF-κB and AP-1. The two NF-κB sites are thought to be essential for HIV-1 transcription (24, 25), whereas the AP-1 sites, while not essential, are thought to enhance HIV-1 transcription (26, 27). Previous studies demonstrated that treatment of HIV-infected MDMs with the TLR2/TLR1 ligand PAM3CSK4 led to an increased association of the p65 subunit of NF-κB and the c-fos subunit of AP-1 with the 5′ LTR which, in turn, correlated with increased virus replication (18); however, the contributions made by each pathway to TLR2-mediated activation have not been previously characterized. To determine the roles of NF-κB and AP-1 in TLR2-activated HIV replication in MDMs, HIV-1-infected cells were treated with either BAY 11-7082, an inhibitor of IκB kinase (28), celastrol, a small molecule inhibitor of the IκB kinase complex (29), or inhibitors that disrupt AP-1 signaling. As shown in Supplemental figure 2, BAY 11-7082 and celastrol treatment completely ablated TLR2/1-enhanced HIV-1 expression. Similarly, the use of an LTR-based reporter construct with mutations in the NF-κB binding sites did not result in increased gene expression in response to TLR2 signaling (Supplemental Figure 2C). Treatment of HIV-infected macrophages with inhibitors of kinases upstream of AP-1 activation, such as MEK1/2 (U0126, PD98509), and p38 (SB203580), resulted in modest, but reproducible, decreases in TLR2-mediated activation of HIV-1 (Supplemental Figure 2D). Similarly, LTR reporter constructs lacking AP-1 binding sites were activated in response to TLR2 signaling at levels similar to that of the WT construct, further demonstrating the non-essential role of AP-1 in TLR2-mediated HIV-1 activation (Supplemental Figure 2E). Although the regulation of HIV-1 transcription through multiple transcription factor binding sites in and adjacent to the 5′ LTR is complex, these data suggest that, in MDMs, TLR2-activated HIV-1 expression is mediated primarily through NF-κB with a minor contribution from AP-1 signaling.

**Figure 2.**
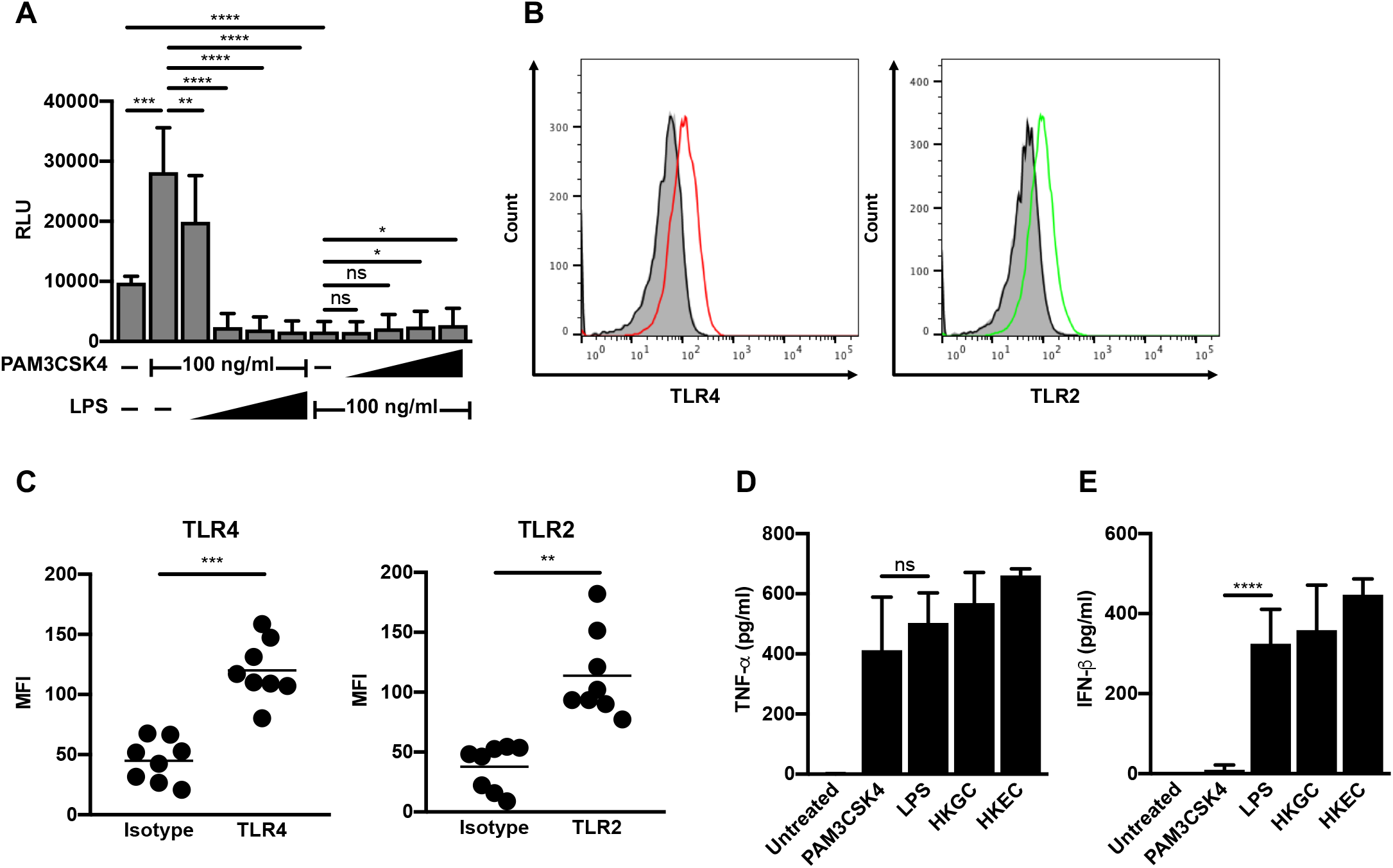
TLR4 signaling is dominant in MDMs. (A) MDMs (2.5 × 10^5^ cells/well) were infected with a single-round, replication-defective HIV-luciferase reporter virus (MOI = 0.1). Forty-eight hours after infection, cells were treated with a fixed concentration of PAM3CSK4 (100 ng/ml) and increasing concentrations of LPS (1-1000 ng/ml, as indicated) or a fixed concentration of LPS (100 ng/ml) and increasing concentrations of PAM3CSK4 (1-1000 ng/ml, as indicated) for 18 hours. Cells were then lysed and assayed for luciferase activity. The data are the mean (± SD) of six donors, each donor tested in triplicate. (B-C) At eight days post-isolation, MDMs were stained with antibodies against TLR2 or TLR4 or relevant isotype controls. Receptor expression was assessed by flow cytometry. Histograms from one representative donor are shown in (B). Grey, unstained cells; black line, isotype control; red line, TLR4; green line, TLR2. Mean fluorescent intensity (MFI) ± SD from eight donors is depicted in (C). (D-E) MDMs (2.5 × 10^5^ cells/well) were treated with the TLR2 ligand PAM3CSK4 (100 ng/ml), the TLR4 ligand LPS (100 ng/ml), heat-killed GC (MOI = 10), or heat-killed *E. coli* (MOI = 10) for 18 hours. Cell supernatant was harvested, filtered through a 0.2 μm filter, and analyzed by ELISA for TNF-α (D) and IFN-β (E) production. Data represent mean (± SD) of seven donors (four donors for heat-killed *E. coli).* *, p < 0.05; **, p < 0.01; ***, p < 0.001; p < 0.0001.

### Co-infection with *Neisseria gonorrhoeae* or *Escherichia coli* represses HIV-1 replication in MDMs

Our preliminary studies using purified TLR ligands in isolation suggested that different TLR signaling cascades had diverse effects on HIV-1 replication. Since most pathogens encode multiple TLR ligands, we sought to determine the effects of intact pathogens on HIV-1 replication. We incubated HIV-infected MDMs with *N. gonorrhoeae* (GC), which expresses ligands for TLR2, TLR4, and TLR9. We found that increasing amounts of GC led to a dosedependent decrease in HIV-1 replication in MDMs (Figure 1F). Bacterial replication was not required for these effects, as heat-killed GC led to repression of HIV-1 replication in MDMs (Supplemental Figure 3A). GC-mediated repression occurred at the level of viral transcription (Figure 1G) and did not decrease viral RNA stability (Supplemental Figure 3B). Similar to what we observed with purified LPS, the biological sex of the donors had no effect on GC- or *E. coli-* mediated HIV-1 repression in MDMs (Supplemental Figure 3C). In addition, repression of HIV-1 replication is not specific for GC, but may be a generalized response to Gram-negative bacteria, as co-infection with *E. coli* also repressed HIV-1 replication in MDMs in a manner similar to GC (Figure 1H). Despite the presence of both activating (TLR2) and repressing (TLR4) TLR ligands, both GC and *E. coli* mediated repression of HIV-1 replication in macrophages. This finding raised several possibilities: 1) the dominance of TLR4 signaling over TLR2 signaling in MDMs; 2) different expression levels of TLR2, TLR4, and TLR4-associated molecules such as CD14 and MD-2 on MDMs; 3) different cytokine profiles produced in response to GC or *E. coli*; and/or 4) variable expression of signaling molecules downstream of TLRs. These scenarios were further explored.

### TLR4 signaling is dominant in MDMs

To determine whether certain TLR pathways are dominant in MDMs, we performed co-treatments of HIV-infected MDMs with the TLR2 ligand PAM3CSK4 and the TLR4 ligand LPS. We found that increasing the concentration of LPS against a fixed concentration of PAM3CSK4 led to a reversal of TLR2-mediated activation of HIV-1 and, eventually, to repression of HIV-1 replication (Figure 2A). Conversely, increasing the concentration of PAM3CSK4 against a fixed concentration of LPS did not reverse LPS-mediated repression of HIV-1 (Figure 2A). Flow cytometry was used to determine that the different responses of MDMs were likely not due to receptor expression, as MDMs express both TLR2 and TLR4 (Figure 2B-C). In addition, MDMs produced both TNF-α and IFN-β in response to LPS treatment, GC co-infection, and *E. coli* co-infection. Whereas treatment of MDMs with LPS resulted in a similar cytokine profile to that of co-infection, treatment of MDMs with the TLR-ligand PAM3CSK4 resulted in the production of TNF-α, but not appreciable levels of IFN-β (Figure 2D-E). Taken together, our data suggest that TLR4 signaling, which negatively regulates LTR-driven gene expression, is dominant in MDMs.

### LPS- and GC-mediated repression of HIV-1 in MDMs is dependent on TRIF-mediated type I IFN production

Since LPS and GC both induce type I IFN production, whereas the TLR2 ligand PAM3CSK4 does not, we wished to determine whether GC-stimulated production of IFN-α/β contributes to repression of HIV-1 in MDMs. We found that treatment of HIV-infected MDMs with the vaccinia virus-encoded soluble type I IFN receptor B18R reversed GC-mediated inhibition of HIV-1 replication, suggesting that TLR4-mediated IFN production is required for HIV-1 repression by GC (Figure 3A). Since both purified TLR4 ligands and GC, which encodes ligands for TLR2, TLR4, and TLR9, repress HIV-1 replication in MDMs, we predicted that downstream effector molecules of TLR4 signaling would contribute to the repression of HIV-1 replication in MDMs. First, we confirmed that TLR4 signaling was responsible for GC-mediated HIV-1 repression in MDMs. Treatment with the TLR4-specific inhibitor TAK242 reversed the LPS- and GC-dependent repression of HIV-1 in MDMs (Figure 3B). Treatment with TAK242 had no effect on TLR2-mediated activation of HIV-1 replication in MDMs, consistent with reports that TAK242 is specific for TLR4 (30).

**Figure 3.**
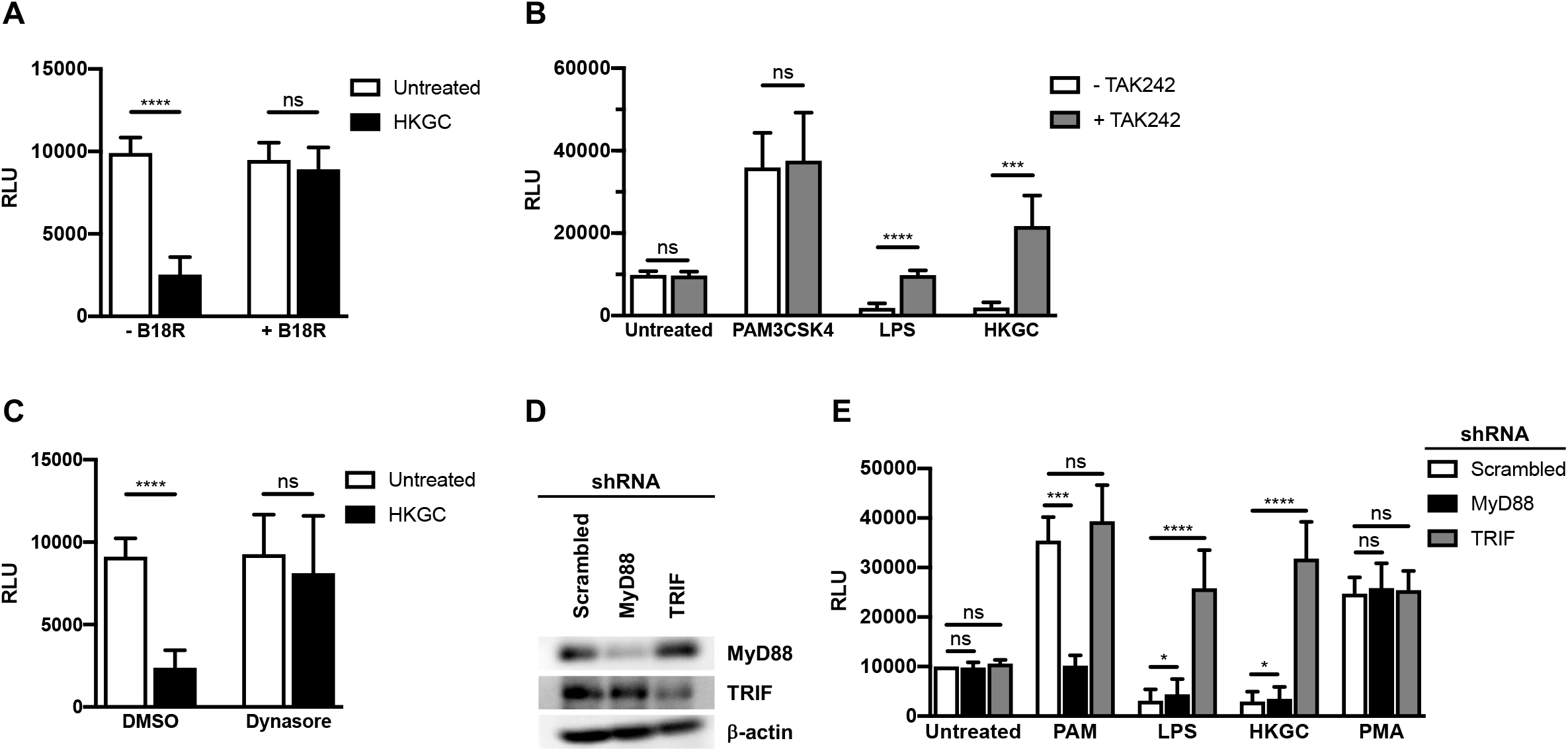
LPS- and GC-mediated repression of HIV-1 replication in MDMs requires TLR4, TRIF, and type I IFNs. (A) MDMs (2.5 × 10^5^ cells/well) were infected with a single-round, replication-defective HIV-luciferase reporter virus (MOI = 0.1). Forty-eight hours after infection, cells were treated with GC (MOI = 10) in the absence (white bars) or presence (black bars) of B18R (100 ng/ml) for 18 hours. The cells were then lysed and assayed for luciferase activity. The data are the mean (± SD) of seven donors, each donor tested in triplicate. (B) MDMs (2.5 × 10^5^ cells/well) were infected as above. Forty-eight hours after infection, cells were treated with PAM3CSK4 (100 ng/ml), LPS (100 ng/ml), or heat-killed GC (MOI = 10) in the absence (white bars) or presence (black bars) of TAK242 (1 μg/ml) for 18 hours. The cells were then lysed and assayed for luciferase activity. The data are the mean (± SD) of six donors, each donor tested in triplicate. (C) MDMs (2.5 × 10^5^ cells/well) were infected as above. Forty-eight hours after infection, cells were treated with vehicle control (white bars) or with (black bars) the dynamin inhibitor Dynasore (80 μM) for 15 minutes prior to treatment with heat-killed GC (MOI = 10) for 18 hours. The cells were then lysed and assayed for luciferase activity. The data are the mean (± SD) of six donors, each donor tested in triplicate. (D-E) MDMs (2×10^6^ cells/well) were transfected with a control scrambled shRNA, shRNA targeting MyD88, or shRNA targeting TRIF. Knock-down of protein expression was detected by western blot (D). Transfected MDMs were infected with a single-round, replication-defective HIV-luciferase reporter virus at an MOI = 0.1. Forty-eight hours after infection, cells were treated with PAM3CSK4 (100 ng/ml), LPS (100 ng/ml), heat-killed GC (MOI = 10), or PMA (10 nM) for 18 hours. The cells were then lysed and assayed for luciferase activity (E). The data are the mean (± SD) of six donors. *, p < 0.05; **, p < 0.01; ***, p < 0.001; p < 0.0001.

It has been shown that TLR4, which can utilize both MyD88 and TRIF adaptor proteins, initiates different signaling pathways dependent upon its cellular location. Cell-surface TLR4 engagement leads to MyD88-dependent signaling, whereas endosomal TLR4 engagement leads to TRIF-dependent signaling (31). To examine whether TRIF-dependent signaling is responsible for HIV-1 repression, we blocked dynamin-dependent endocytosis of TLR4 with Dynasore, which prevents TRIF-dependent signaling while leaving MyD88-dependent signaling intact. As shown, blocking endocytosis-mediated TLR4 internalization (Supplemental figure 4) reversed GC-mediated repression of HIV-1 in MDMs (Figure 3C). Given the ability of GC to signal through both TLR2-MyD88 and TLR4-TRIF, one might expect the inhibition of type I IFN signaling by B18R or the inhibition of endocytosis by Dynasore to lead to augmented viral gene expression through intact MyD88 signaling. However, we did not observe this, likely due to incomplete inhibition of either IFN signaling or endocytosis.

To confirm the role of MyD88 in TLR2-mediated HIV-1 activation and TRIF in TLR4-mediated HIV-1 repression, we used shRNAs to knock down the two molecules in HIV-infected MDMs (Figure 3D). Knock-down of MyD88 led to a loss of TLR2-mediated HIV-1 activation, but had no effect on LPS or GC-mediated HIV-1 repression (Figure 3E). In contrast, knock-down of TRIF had no effect on TLR2-mediated HIV-1 activation, but reversed LPS- and GC-mediated repression of HIV-1 replication (Figure 3E). Knock-down of either MyD88 or TRIF had no effect on the activation of HIV-1 by the phorbol ester PMA, which signals directly through protein kinase C, independently of TLRs (Figure 3E). These data suggest that the TLR4-TRIF-type I IFN axis in MDMs leads to GC- and *E. coli*-mediated repression of HIV-1 replication.

### TLR4 signaling leads to differential IRF recruitment to the HIV-1 LTR

Since type I IFN production is critical for GC- and *E. coli*-mediated HIV-1 repression in MDMs, we examined the role of interferon-stimulated genes (ISGs) in HIV-1 regulation. Previous studies have shown that ISGs are temporally regulated in macrophages in response to innate immune sensors and type I IFN signaling (32, 33). To determine whether the repressive effects of LPS were due to early or late phase ISGs, HIV-1-infected MDMs were treated with the TLR2 ligand PAM3CSK4 or the TLR4 ligand LPS and total cytoplasmic RNA was extracted at various times post treatment. Treatment of HIV-infected MDMs with the TLR2 ligand PAM3CSK led to a continuous increase in HIV-1 RNA levels (Figure 4A). In contrast, treatment of HIV-infected MDMs with the TLR4 ligand LPS led to an initial short-lived increase in HIV-1 RNA levels; however, levels steadily declined thereafter (Figure 4A), indicating that HIV-1 transcription displays a biphasic response to TLR4 stimulation in MDMs. This suggests that late-phase proteins induced by type I IFNs are responsible for TLR4-mediated decreases in HIV-1 transcription. It is known that HIV-1 contains an interferon-stimulated response element (ISRE) in the Gag-leader sequence (GLS), immediately downstream from the 5′ LTR. Because type I IFN is required for LPS- and GC-mediated repression of HIV-1 in MDMs, we assessed the role of the ISRE in this process using transient transfection assays with mutated LTR-reporter constructs in HEK293 cells expressing TLR4, MD-2, and CD14. We found that LPS treatment repressed LTR-driven reporter-gene expression in cells expressing WT ISRE elements, but not in cells transfected with an LTR-luciferase construct containing a mutated ISRE (Supplemental Figure 5). This suggests that transcription factor engagement of the ISRE governs TLR4-mediated HIV-1 repression.

**Figure 4.**
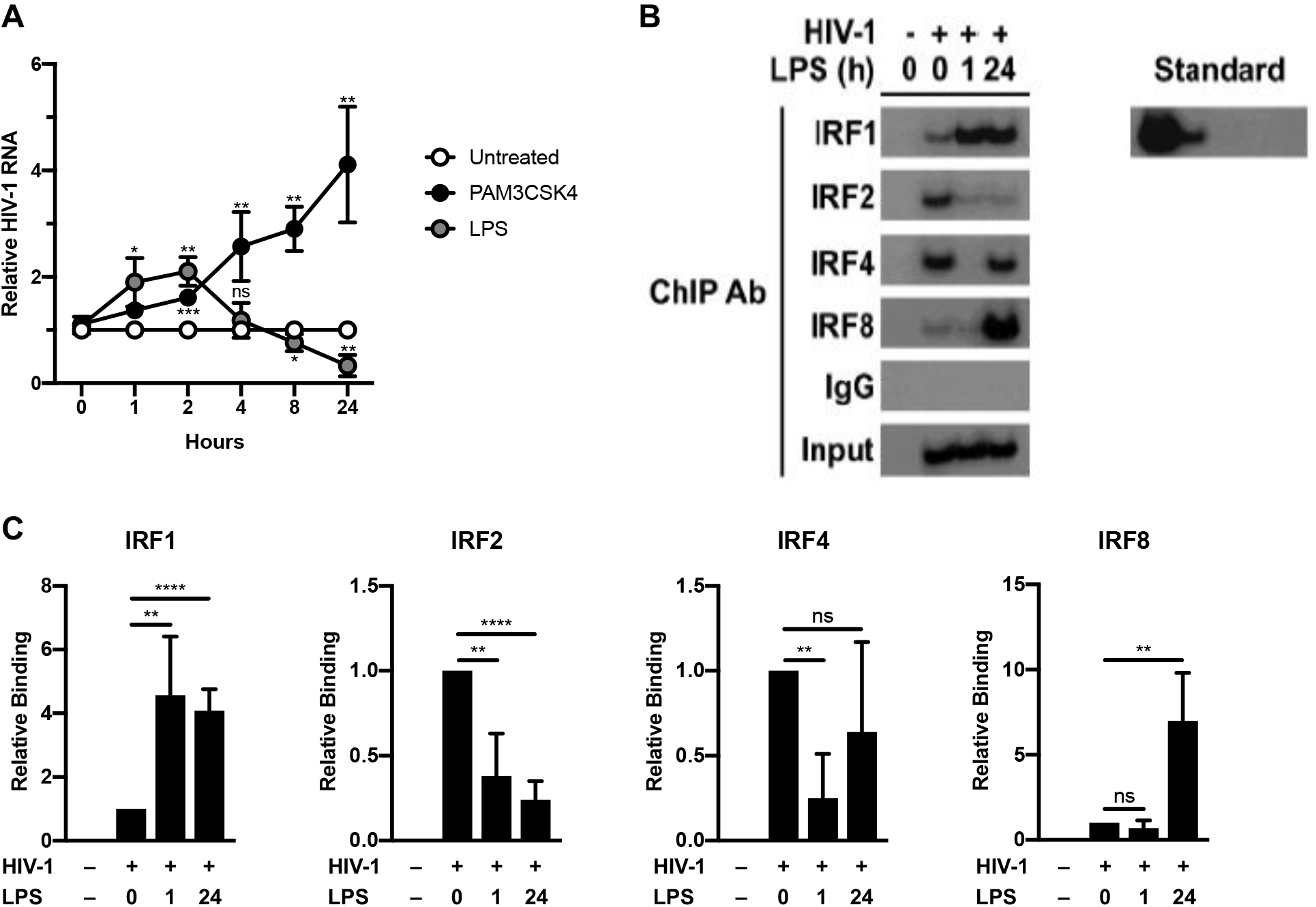
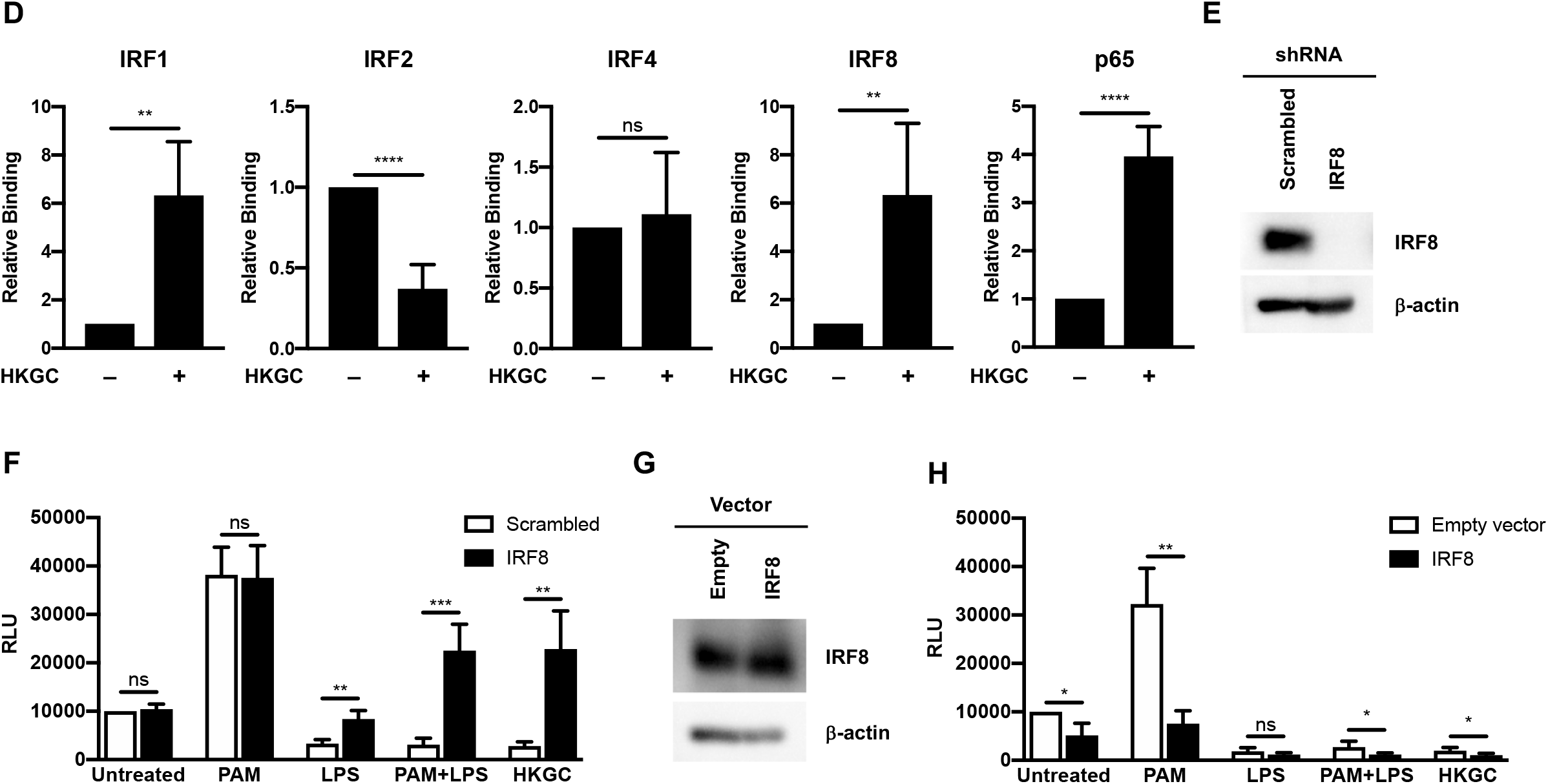
LPS- and GC-mediated repression of HIV-1 in MDMs is associated with changes in IRF recruitment to the ISRE. (A) MDMs (2×10^6^ cells/well) were infected with a single-round, replication-defective HIV-luciferase reporter virus (MOI = 0.1). Forty-eight hours after infection, cells were treated with PAM3CSK4 (100 ng/ml) or LPS (100 ng/ml). At various time points after TLR stimulation, cells were harvested, lysed, and total cytoplasmic RNA was extracted. Viral RNA accumulation was assessed by RT-PCR. The data are the mean (± SD) of four donors. (B-C) MDMs (1.2 × 10^7^ cells/plate) were infected with a single-round replication-defective HIV-GFP reporter virus at an MOI = 2. Forty-eight hours after infection, cells were treated with LPS (100 ng/ml). At either 1 or 24 hours after LPS treatment, cells were fixed with formaldehyde, lysed, sonicated, and subjected to immunoprecipitation with antibodies against IRF1, IRF2, IRF4, IRF8, or rabbit IgG (isotype control). Association with the HIV-1 ISRE was assessed by PCR using HIV-1 specific primers. Data from one representative donor is depicted in (B). Composite data representing the mean (± SD) from five donors are showin in (C). (D) MDMs (1.2 × 10^7^ cells/plate) were infected with a single-round replication-defective HIV-GFP reporter virus at an MOI = 2. Forty-eight hours after infection, cells were treated with heat-killed GC (MOI = 10). Twenty-four hours after GC treatment, cells were fixed with formaldehyde, lysed, sonicated, and subjected to immunoprecipitation with antibodies against IRF1, IRF2, IRF4, IRF8, or rabbit IgG (isotype control). Association with the HIV-1 ISRE was assessed by PCR using HIV-1 specific primers. Composite data from five donors are shown. (E-F) MDMs (2×10^6^ cells/well) transfected with a controlled scrambled shRNA (white bars) or with shRNA targeting IRF8 (black bars) were infected with a single-round, replication-defective HIV-luciferase reporter virus at an MOI = 0.1. Knock-down of protein expression was detected by western blot (F). Forty-eight hours after infection, cells were treated with PAM3CSK4 (100 ng/ml), LPS (100 ng/ml, a combination of PAM3CSK4 and LPS (each at 100 ng/ml), or GC (MOI = 10) for 18 hours. Cells were then lysed and assayed for luciferase activity (F). The experiment was performed using cells from five different donors. (G-H) MDMs transfected with an empty vector (white bars) or a vector encoding IRF8 (black bars) were infected with a single-round, replication-defective HIV-luciferase reporter virus at an MOI = 0.1. IRF8 protein expression was detected by western blot (G). Forty-eight hours after infection, cells were treated with PAM3CSK4 (100 ng/ml), LPS (100 ng/ml, a combination of PAM3CSK4 and LPS (each at 100 ng/ml), or GC (MOI = 10) for 18 hours. Cells were then lysed and assayed for luciferase activity (H). The experiment was performed using cells from four different donors. *, p < 0.05; **, p < 0.01; ***, p < 0.001; p < 0.0001.

Previous studies have shown that IRF1 and IRF2 both bind to this ISRE *in vitro* and that IRF1 and IRF2 expression are associated with enhanced HIV-1 transcription (34). Two other IRFs, IRF4 and IRF8, are also expressed in macrophages (35) and have been shown to increase in levels in response to type I IFN signaling and other signals (36, 37). Interestingly, IRF8 has been implicated in maintaining HIV-1 latency in infected monocytic cell lines (34, 38, 39), and IRF4 has been implicated in negative regulation of TLR signaling (40). We therefore investigated whether various IRFs are recruited to the HIV-1 ISRE in response to LPS and GC treatment in MDMs. Using chromatin immunoprecipitation analysis, we found that IRF1, IRF2, IRF4, and IRF8 all are able to associate with the 5′ LTR and GLS containing the ISRE in HIV-infected MDMs (Figure 4C-D). Early after treatment with LPS, the levels of IRF1 associated with this region of the viral promoter increased, whereas the levels of IRF2 and IRF4 decreased. By 24h post-treatment with LPS, the levels of IRF4 and IRF8 associated with this region increased. Of particular note, the levels of IRF8 recruitment increased well above those seen in unstimulated MDMs (Figure 4C). A similar pattern of IRF recruitment to the 5′ LTR and GLS occurred in GC-treated MDMs (4D), suggesting that repression of HIV-1 transcription in response to LPS and GC treatment is due to enhanced IRF8 recruitment to the 5′ LTR and GLS. To confirm the central role of IRF8 in TLR4-mediated repression of HIV-1 expression in MDMs, we used shRNA to knockdown IRF8 expression in MDMs (Figure 4E). Reducing IRF8 expression reversed TLR4-mediated HIV-1 repression in response to treatment with LPS or GC. Knockdown of IRF8 led to activation of HIV-1 expression in cells treated with a combination of PAM3CSK4 and LPS or GC, similar to that seen with treatment with PAM3CSK4 alone (Figure 4F). In contrast, overexpression of IRF8 in MDMs led to decrerased HIV-1 expression in untreated MDMs and reversed the activation of HIV-1 expression in PAM3CSK4-treated MDMs, but had no effect on LPS-mediated repression in MDMs (Figure 4G-H). There was a small, but significant, enhancement of HIV-1 repression in MDMs treated with a combination of PAM3CSK4 and LPS or with GC.

Taken together, these data suggest that both LPS and GC activate TLR4-mediated TRIF signaling in MDMs, resulting in the production of type I IFNs. In turn, IFNs work in an autocrine or paracrine fashion to induce the expression of IRF8, which then binds to the ISRE present in the GLS of HIV-1 to repress viral transcription (Figure 6).

### LPS and GC-treatment induces persistent low-level/latent HIV-1 infection in MDMs

Recent studies in animals and human tissues demonstrate that HIV-1 can form persistent low-level or latent infections in macrophages (41–44). Our data suggest that engagement of the TLR4-TRIF-type I IFN axis in macrophages can repress virus replication and we wished to determine whether signaling through this axis could contribute to the establishment of persistent low-level or latent HIV-1 infection in macrophages. To this end, HIV-1-infected MDMs were treated a single time with the TLR2 ligand PAM3CSK4, the TLR4 ligand LPS, heat-killed GC, IFN-α, or IFN-β at day 3 post-infection. As shown in Figure 5, while there was a range of virus replication in the various donors, we found that treatment with a single dose of LPS, heat-killed GC, IFN-α, or IFN-β consistently led to a prominent, sustained decrease in HIV-1 replication in MDMs, whereas treatment with PAM3CSK4 led to a transient increase in HIV-1 replication followed by a slight decrease in replication. These data suggest that engagement of the TLR4-TRIF-type I IFN axis can promote low-level persistent/latent HIV-1 infection in MDMs.

**Figure 5.**
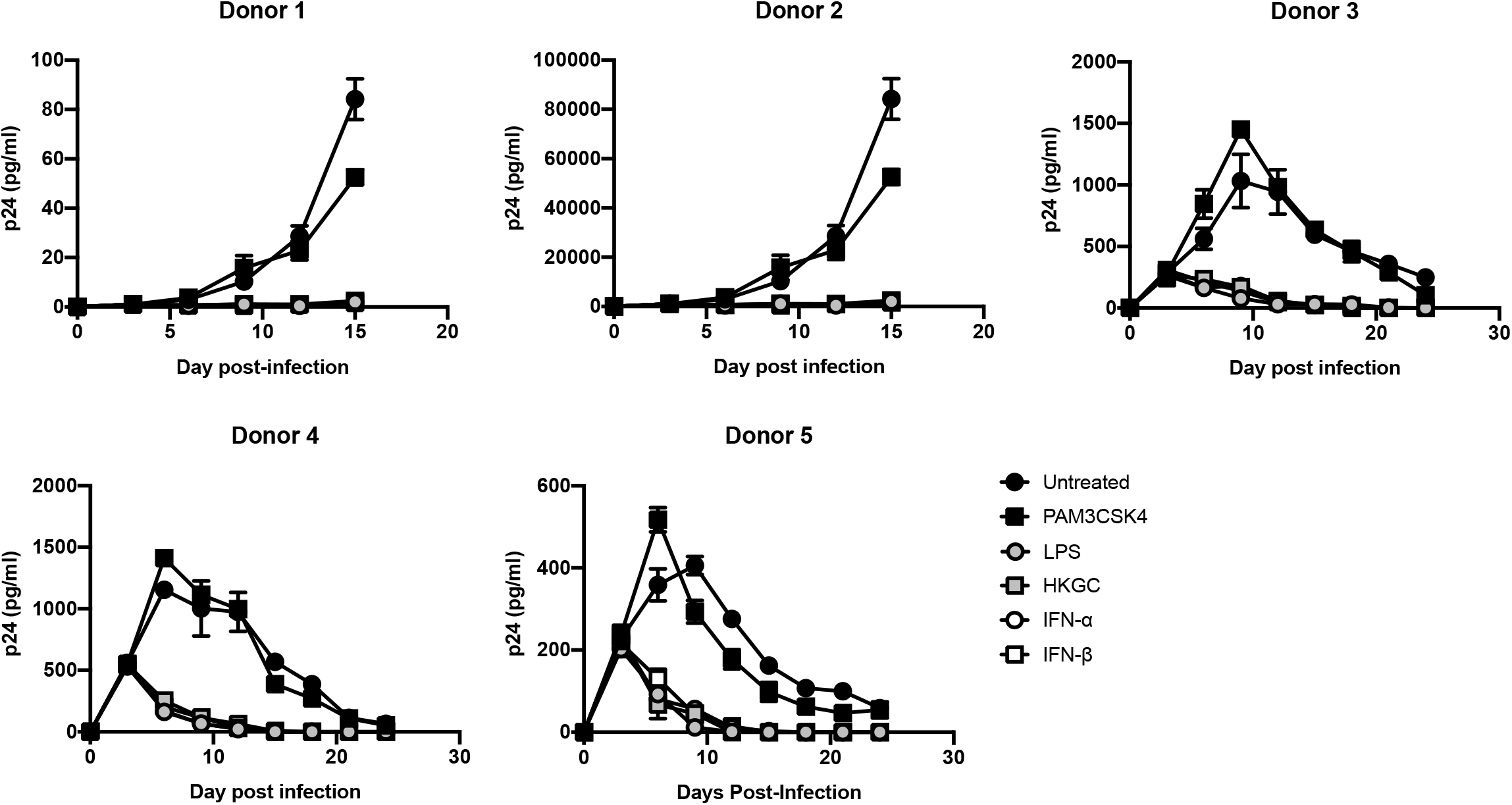
Treatment with LPS, GC, or type I IFNs induces a low-level persistent/latent infection in MDMs. MDMs (2.5×10^5^ cells/well) were infected with replication-competent HIV-1_Ba-L_ (MOI of 0.1). At day 3 post-infection, the cells were treated with a single dose of PAM3CSK4 (100 ng/ml), LPS (100 ng/ml), GC (MOI = 10), IFN-α (1000 U/ml), or IFN-β (1000 U/ml). Cell-free supernatants were harvested every three days, and virus production was monitored by p24 ELISA. Data from five independent donors, each tested in triplicate, are shown. *, p < 0.05; **, p < 0.01; ***, p < 0.001; p < 0.0001.

## DISCUSSION

In these studies, we provide evidence that the interaction between commensal and pathogenic bacteria can repress HIV-1 replication in macrophages by altering the recruitment of transcription factors to the HIV-1 GLS, thereby inducing a state of proviral latency. We further demonstrate that TLR2 ligands trigger MyD88-mediated signaling that increases virus expression via the activation of NF-κB, whereas TLR4 ligands trigger TRIF-dependent production of type I IFNs. Type I IFN signaling, in turn, is associated with the recruitment of IRF8 to the ISRE located in the GLS and a shift to low-level or latent HIV-1 infection.

A number of studies have shown that IRFs play an important role in the regulation of HIV-1 replication. There is an ISRE located downstream from the 5′ LTR in the GLS that is essential for efficient viral replication (26, 45). This ISRE is typically bound by IRF1 and/or IRF2, leading to activation of virus transcription (34, 46) through the recruitment of transcriptional coactivators, such as the histone acetyl transferase (HAT) p300/CBP (47). IRF1 and IRF2 are ubiquitously expressed in cells, thought they can be upregulated by type I IFNs (36) and, in the case of IRF1, by TLR signaling (48, 49) and HIV-1 infection (45, 50), illustrating how HIV-1 can co-opt the antiviral IFN response to augment its own replication. Once associated with the

ISRE, IRF1 can cooperatively bind to both NF-κB at the HIV-1 LTR and the viral transactivator Tat at the HIV-1 TAR loop to augment viral transcription/elongation (34, 51). Our studies demonstrate that both IRF1 and IRF2 associate with the HIV-1 ISRE in unstimulated MDMs (Fig. 4). Upon stimulation with TLR4 ligands, IRF1recruitment to the HIV-1 ISRE is enhanced (Fig. 4), consistent with the prevailing theory that TLR-MyD88 signaling can activate IRF1 (52). This is accompanied by a concomitant decrease of IRF2 binding. These data suggest that IRF1 binding to the ISRE as either monomers or homodimers activates HIV-1 expression, whereas IRF2 binding to the ISRE as monomers, homodimers, or heterodimers with IRF1 represses HIV-1 expression. Unfortunately, ChIP analysis of HIV-infected MDMs using current tools does not permit differentiating between the association of various homodimers and heterodimers with the ISRE at a population level.

We demonstrate that at late time points after TLR4 engagement, IRF8 is recruited the the GLS downstream from the 5′ LTR (Figure 4), and that this is associated with decreased HIV-1 transcription (Figure 1). Macrophages express high basal levels of IRF8, although its expression can be further enhanced in response to type I IFNs (36, 37) or TLR signaling (53, 54). IRF8 has been shown to bind to IRF1, in addition to other transcription factors, and to serve as either a transcriptional activator or a transcriptional inhibitor of other genes in a context-dependent manner (55–57). Previous studies have shown that IRF8 can repress HIV-1 expression (34, 38, 39). In fact, the interaction between IRF8 and IRF1 has been shown to repress HIV-1 transcription in Jurkat cells (34). This may be due to IRF8-mediated disruption of the IRF1-Tat interaction and/or the IRF1-NF-κB interaction (51) that increase viral replication. Based on our data, we propose that changes in the IRF binding pattern to the ISRE in response to TLR signaling have profound effects on HIV-1 replication. In unstimulated HIV-infected macrophages, the ISRE is most likely occupied by IRF1/IRF2 heterodimers that allow for a low-level of virus replication (Figure 6A). Early after stimulation of TLR4 with LPS, there is a switch to IRF1 homodimers present at the ISRE that allow for high levels of virus replication due to cooperative binding between IRF1, NF-κB, and HIV-1 Tat. (Figure 6B). At late time points after TLR4 stimulation with LPS or GC, during the IFN feedback phase of the response, the ISRE is occupied by IRF1/IRF8 heterodimers (Figure 6C). These IRF1/IRF8 heterodimers likely block the cooperative interaction(s) between IRF1, NF-κB, and Tat, thereby repressing HIV-1 replication. Although we also demonstrate that IRF4 is recruited transiently to the HIV-1 ISRE following treatment with LPS, the biological significance of this finding is uncertain. Prior studies have provided evidence for an LPS/TLR4-mediated repression of HIV-1 expression through the induction of type I IFNs and other mechanisms (16, 17, 58–62). Our data extend these findings and demonstrate that LPS treatment, as well as infection with the sexually transmitted pathogen GC or the gut-associated microbe *E. coli,* represses HIV-1 expression in MDMs through the TLR4-mediated, TRIF-dependent production of type I IFNs and the subsequent recruitment of IRF8 to the HIV-1 ISRE.

**Figure 6.**
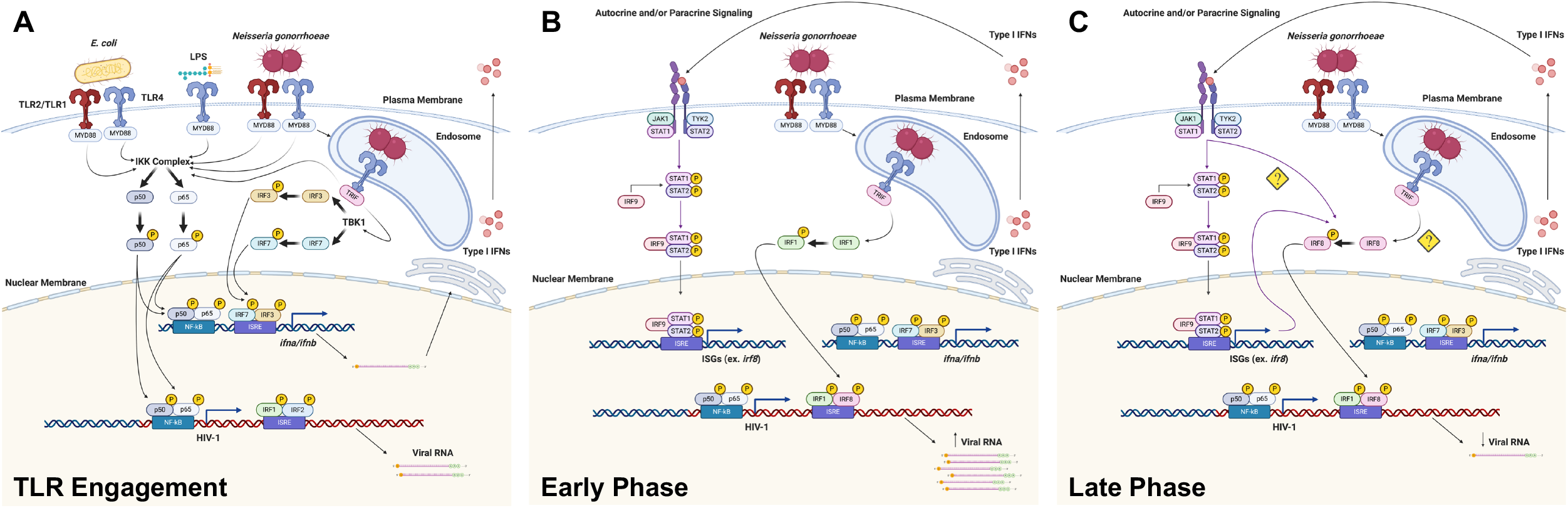
Co-infection with GC or *E. coli* represses HIV-1 replication by altering IRF recruitment to the HIV-1 GLS. (A) Upon engagement of TLR2 and/or TLR4 by GC or *E. coli* at the cell surface, transcription factors such as NF-κB are phosphorylated and are subsequently recruited the HIV-1 5′ LTR to drive viral transcription in MDMs. Upon engagement of TLR4 in the endosome, IRF3 and IRF7 are phosphorylated and recruited to the IFN-α and/or IFN-β promoters to drive type I IFN expression. (B) During the early phase of the response, type I IFNs drive the expression of ISGs, including IRFs. Signaling through endosomal TLR4 leads to the activation of IRF1 and its recruitment to the HIV-1 GLS, further enhancing HIV-1 transcription. (C) During the late phase of the response there is a type I IFN-dependent activation of IRF8 and subsequent recruitment to the HIV-1 GLS, thereby repressing HIV-1 transcription.

We have demonstrated that MyD88-dependent signaling activates HIV-1 expression in MDMs by activating NF-κB, whereas TRIF-dependent signaling represses HIV-1 expression in these cell types through the type I IFN-mediated recruitment of IRF8 to the HIV-1 ISRE. We have also shown that TLR4-mediated, TRIF-dependent signaling is dominant in MDMs, both in the context of treatment with purified TLR ligands as well as infection with the sexually transmitted pathogen GC or the gut-associated microbe *E. coli.* Co-infection of virus-infected MDMs by pathogens that express TLR4 ligands decreases HIV-1 replication through the recruitment of IRF8 to the HIV-1 ISRE. Prior studies have provided evidence for an LPS/TLR4-mediated repression of HIV-1 expression through the induction of type I IFNs and other mechanisms. These studies employed HIV-1 infection of myeloid cells (16, 17, 58–61) or transfection using full-length viral constructs (62). Our data extends these findings and demonstrates that LPS treatment represses HIV-1 expression in MDMs through its interaction with TLR4. Specifically, we show that LPS induces TLR4-mediated, TRIF-dependent production of type I IFNs, which in turn lead to the repression of HIV-1 replication in MDMs through the recruitment of IRF8 to the HIV-1 ISRE.

Importantly, our data suggest that the microbial environment can influence the state of HIV-1 replication and the establishment of latency in human macrophages as part of the viral reservoir in infected individuals under antiretroviral therapy (ART) regimens. Macrophages can be productively infected with HIV-1 *in vivo* and viral replication can be modulated by co-pathogens through their interactions with innate immune receptors such as TLRs (18, 63). We demonstrate that productive infection of macrophages can be altered by TLR signaling in response to purified ligands and bacterial co-infection, with TLR2- and TLR5-mediated signaling activating HIV-1 and TLR3- and TLR4-mediated signaling repressing HIV-1 replication in MDMs (Figure 1).

In addition to their role in HIV-1 production, macrophages also contribute to HIV-1 persistence *in vivo.* Although CD4+ memory T cells are thought to constitute the majority of the HIV-1 reservoir, several studies have demonstrated that tissue resident macrophages in the lymph nodes (64–66), gastrointestinal tract (5, 67), genitourinary tract (2, 42, 68), liver (69–71), and lung (72–74), as well as perivascular macrophages and microglial cells in the brain (41,75–80), can serve as tissue reservoirs for HIV-1. In SHIV-infected rhesus macaques, *in vivo* viral replication was sustained by tissue macrophages after depletion of CD4+ T cells (81). Moreover, HIV-1 persistence in macrophages was confirmed in HIV-1 infected humanized myeloid-only mice in which viral rebound was observed in a subset of the animals following treatment interruption (43). These studies demonstrate that macrophages have the capacity to serve as *bona fide* HIV-1 reservoirs *in vivo.* Our findings that pathogenic and commensal bacteria, through engagement of TLRs, can influence HIV-1 replication in macrophages have potential clinical significance. For example, sexually transmitted infections (STIs) that induce robust type interferon production, such as GC or HSV-2, may repress virus replication in genitourinary tract macrophages that harbor HIV-1 provirus and contribute to viral escape from the immune system and from ART.

The major obstacle to the eradication of HIV-1 is the presence of a persistent viral reservoir that that can resurface upon discontinuation of ART. The potential contribution of HIV-1 in tissue macrophages to virus rebound with the cessation of ART is not entirely understood, but recent primate studies suggest that the functional macrophage reservoir can contribute to viral rebound upon treatment cessation (82–84). Our data demonstrate that interactions between macrophages and pathogenic or commensal microorganisms within the genitourinary and gastrointestinal tracts, such as GC and *E. coli,* may alter the ability of macrophages to serve as reservoirs for viral persistence in the host. Our findings are consistent with independent studies that demonstrate that repeated stimulation of M1-polarized MDMs with proinflammatory cytokines (TNF-α) and/or type II IFNs (interferon-γ) induce a state akin to HIV-1 latency (85). In addition, the oral pathogen *Porphyromonas gingivalis* has been shown to influence the establishment and maintenance of persistent HIV-1 infection in MDMs (86). Finally, studies have demonstrated that a subset of HIV-1-infected macrophages enter a state of viral latency characterized by altered metabolic signatures (87) and apoptotic mechanisms (87). Taken together, these studies demonstrate that co-infection, inflammatory stimuli, and metabolic alterations can influence the establishment and maintenance of the HIV-1 reservoir in macrophages. As an example, gastrointestinal macrophages constitute a major cellular reservoir for HIV-1 (5, 88–90) and are frequently exposed to microbes and microbial products either through luminal sampling (91) or microbial translocation, the latter of which is increased in HIV-positive individuals (92). Our data suggest that interactions such as those between intestinal macrophages and gut-associated microbes may have clinical significance for the establishment and maintenance of the latent HIV-1 reservoir.

Our results demonstrating that Neisseria gonorrhoeae and E. coli repress HIV-1 replication in macrophages by altering transcription factor recruitment to the HIV-1 GLS and induce a state of viral latency confirm the need for further in vitro, ex vivo, and in vivo studies regarding the effects of sexually-transmitted pathogens and commensal microbes on HIV-1 persistence.

## ACKNOWLEDGEMENTS

The authors gratefully acknowledge Dr. Laura Dickey for her scientific and editorial suggestions. The following reagents were obtained through the AIDS Research and Reference Reagent Program, Division of AIDS, NIAID, NIH: HIV IG from NABI and National Heart Lung and Blood Institute (Dr. Luiz Barbosa); anti-p24gag monoclonal antibody clone 183-H12-5C from Dr. Bruce Chesebro and Kathy Wehrly; and MAGI-CCR5 cells from Dr. Julie Overbaugh. The model figure was created with Biorender.com.

This work was supported by funds obtained from the National Institutes of Health (www.nih.gov) grants AI073149, (G.V.), AI143567-02 (V.P.), T32-AI07309 (T.M.H.), and T32-AI0764206 (T.M.H.), a Boston University Division of Graduate Medical Sciences Graduate Student Research Fellowship award (T.M.H.), and a Resident Research Award from the Department of Pathology at the University of Utah (T.M.H.).

## AUTHOR CONTRIBUTIONS

T.M.H. and G.A.V. designed the study. T.M.H and G.A.V. developed the methodology. T.M.H. conducted experiments. T.M.H., V.P., and G.A.V. wrote the manuscript. T.M.H., V.P., and G.A.V. acquired funds. T.M.H. and G.A.V. supervised the study.

---

**Supplemental Figure 1.**
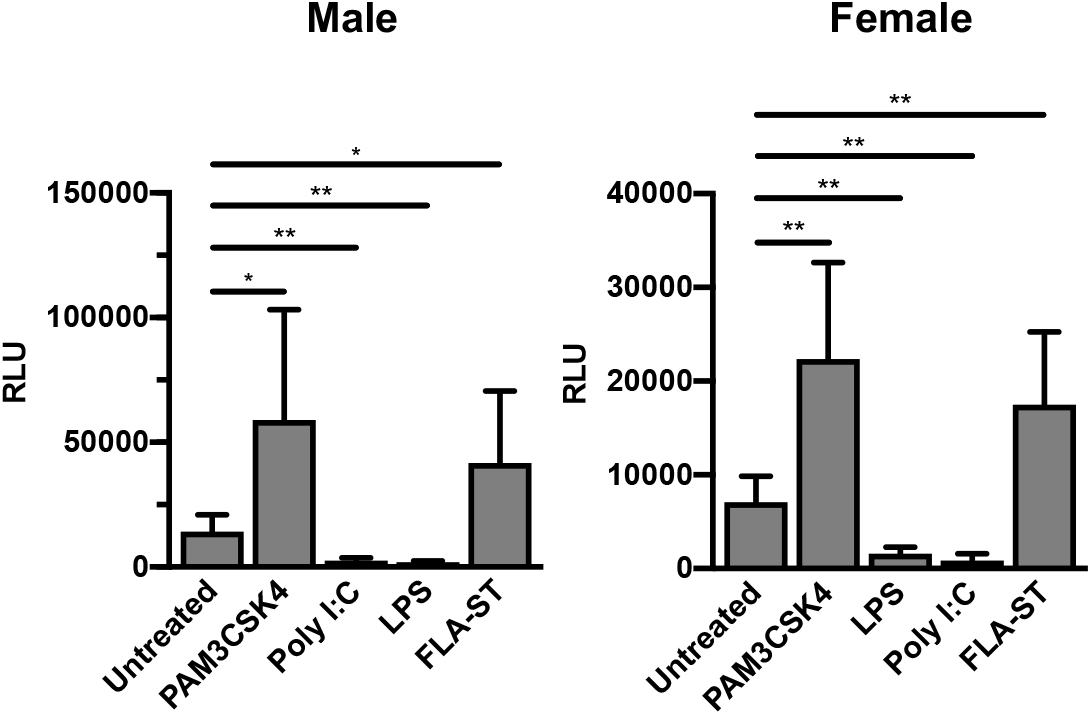
Altered HIV-1 expression in MDMs in response to TLR engagement is not sex-dependent. MDMs (2.5 × 10^5^ cells/well) were infected with a single-round, replication-defective HIV-luciferase reporter virus (MOI = 0.1). After four hours, unbound virus was removed by washing with PBS and cells were cultured in complete medium. Forty-eight hours after infection, cells were treated with the TLR2 ligand PAM3CSK4 (100 ng/ml), the TLR3 ligand poly I:C (25 μg/ml), the TLR4 ligand LPS (100 ng/ml), or the TLR5 ligand FLA-ST (100 ng/ml) for 18 hours. The cells were then lysed and assayed for luciferase activity. Bars represent the mean (± SD) of five male donors and five female donors; each donor tested in triplicate. Although virus replication was decreased overall in MDMs from female donors compared to MDMs from male donors, it was activated by treatment with PAM3CSK4 and FLA-ST, and repressed by Poly I:C and LPS, in a manner similar to that seen in MDMs from male donors. *, p < 0.05; **, p < 0.01; ***, p < 0.001; p < 0.0001.

**Supplemental Figure 2.**
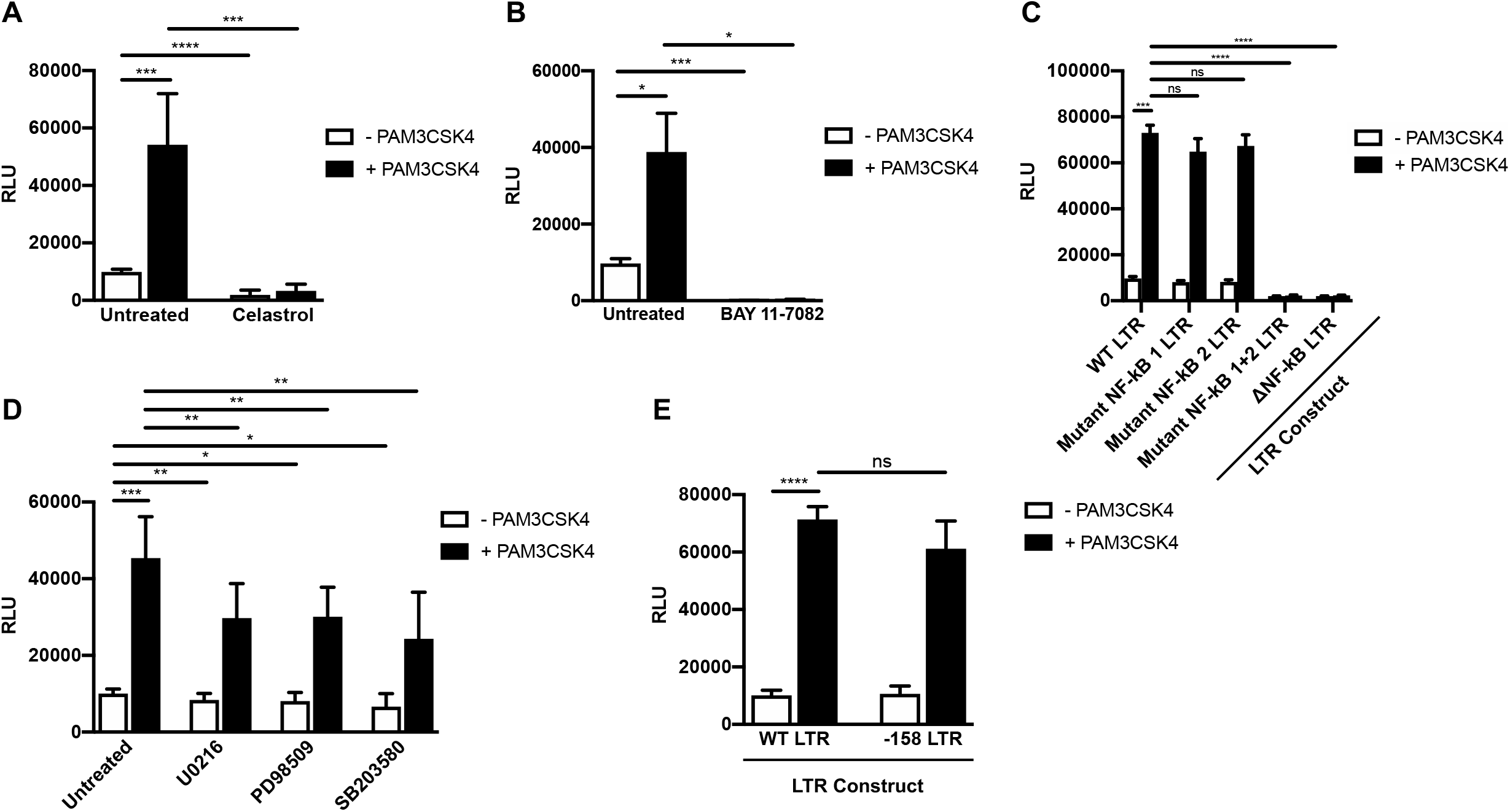
TLR2-activated HIV-1 expression is mediated primarily through NF-κB. (A-B) MDMs (2.5 × 10^5^ cells/well) were infected with a single-round, replication-defective HIV-luciferase reporter virus (MOI = 0.1). Forty-eight hours after infection, cells were treated with PAM3CSK4 (100 ng/ml) in the presence or absence of 10 μM celastrol (A) or 10 μM BAY 11-7082 (B) for 18 hours. The cells were then lysed and assayed for luciferase activity. The data are the mean (± SD) of six donors (celastrol) or three donors (BAY 11-7082); each donor tested in triplicate. (C) HEK293-TLR2^CFP^TLR1^YFP^ cells were transfected with HIV-1 LTR-luciferase reporter constructs with intact NF-κB, mutated NF-κB, or deleted NF-κB binding sites. Following transfection, cells were treated with PAM3CSK4 (100 ng/ml) for 18 hours and then harvested and assayed for luciferase activity. Data represent the mean (± SD) of three independent experiments, each performed in triplicate. (D) MDMs (2.5 × 10^5^ cells/well) were infected as in (A). Forty-eight hours after infection, cells were treated with PAM3CSK4 (100 ng/ml) in the presence or absence of U0126 (10 μM), PD98059 (50 μM), or SB203580 (10 μM) for 18 hours. Cells were then lysed and assayed for luciferase activity. The data are the mean (± SD) of six donors; each donor tested in triplicate. (E) HEK293-TLR2^CFP^TLR1^YFP^ cells were transfected with HIV-1 LTR-luciferase reporter constructs with intact AP-1 sites (WT LTR) or deleted AP-1 binding sites (−158 LTR). Following transfection, cells were treated with PAM3CSK4 (100 ng/ml) for 18 hours and then harvested and assayed for luciferase activity. Data are the mean (± SD) of three independent experiments, each performed in triplicate. *, p < 0.05; **, p < 0.01; ***, p < 0.001; p < 0.0001.

**Supplemental Figure 3.**
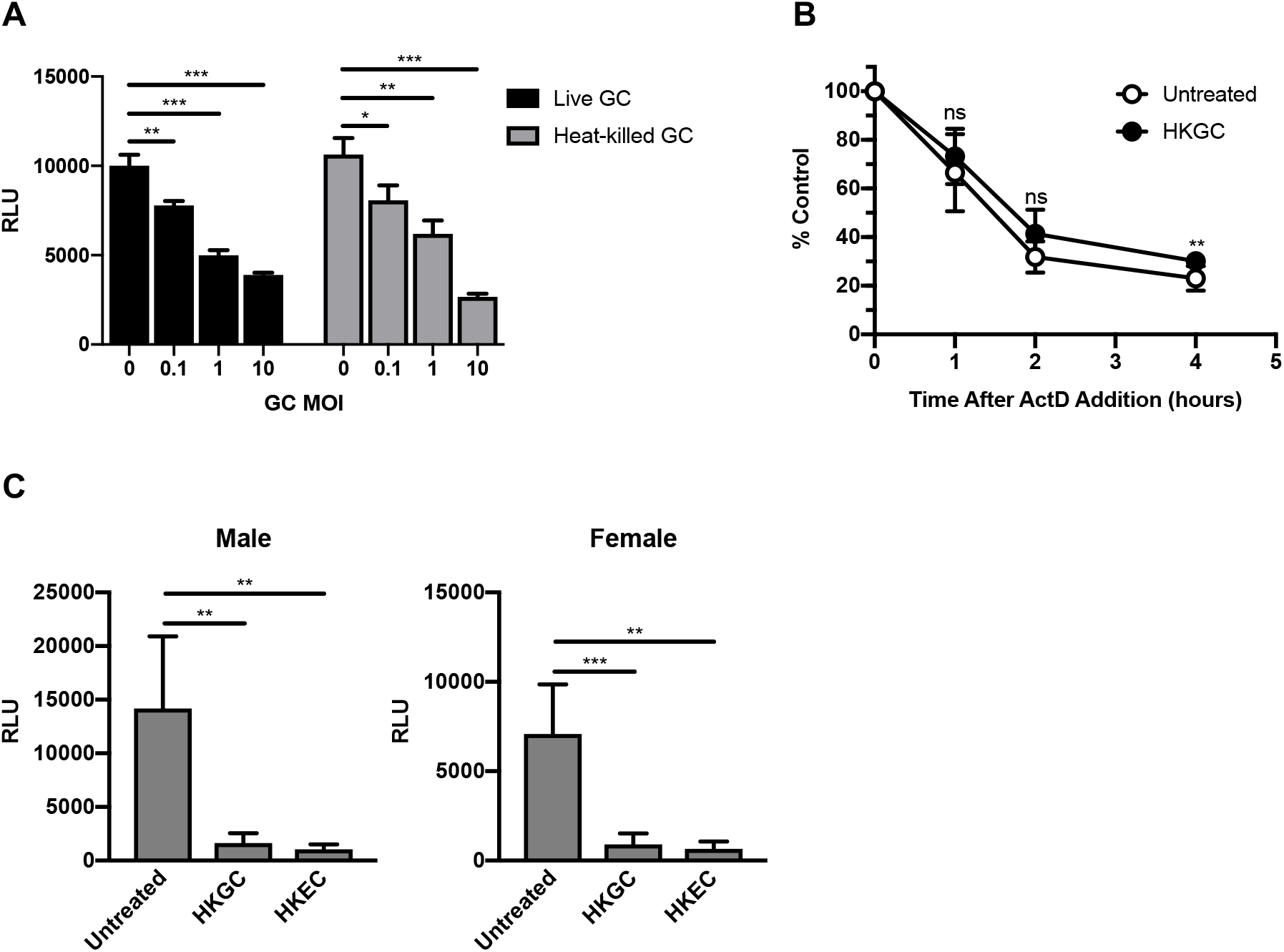
Heat-killed GC represses HIV-1 replication in MDMs, but activates it in MDDCs. (A) MDMs were infected with a single-round, replication-defective HIV-1 luciferase reporter virus (MOI = 0.1). Forty-eight hours after infection, the cells were cultured with increasing amounts of live or heat-killed (56°C treatment) GC overnight. The cells were then lysed and luciferase activity was measured. The data are the mean (± SD) of three donors, each donor tested in triplicate. (B) MDMs (1 × 10^6^ cells/well) were infected as in (A). Forty-eight hours after infection, cells were treated with heat-killed GC (MOI = 10) for 4 h. Cells were then treated with actinomycin D (10 μg/ml) to inhibit transcription. Total cytoplasmic RNA was prepared from the treated cultures at the indicated time points following actinomycin D treatment and analyzed by RT-qPCR for the expression of HIV-1 RNA. The data are the means (± SD) from four donors. (C) MDMs were infected as in (A). Forty-eight hours after infection, cells were treated with heat-killed GC (MOI = 10) or heat-killed *E. coli* (MOI = 10) for 18 hours. The cells were then lysed and assayed for luciferase activity. Bars represent the mean (± SD) of five male donors and five female donors; each donor tested in triplicate. Although virus replication was decreased overall in MDMs from female donors compared to MDMs from male donors, it was repressed by HKGC and HKEC in a manner similar to that seen in MDMs from male donors. *, p < 0.05; **, p < 0.01; ***, p < 0.001; p < 0.0001.

**Supplemental Figure 4.**
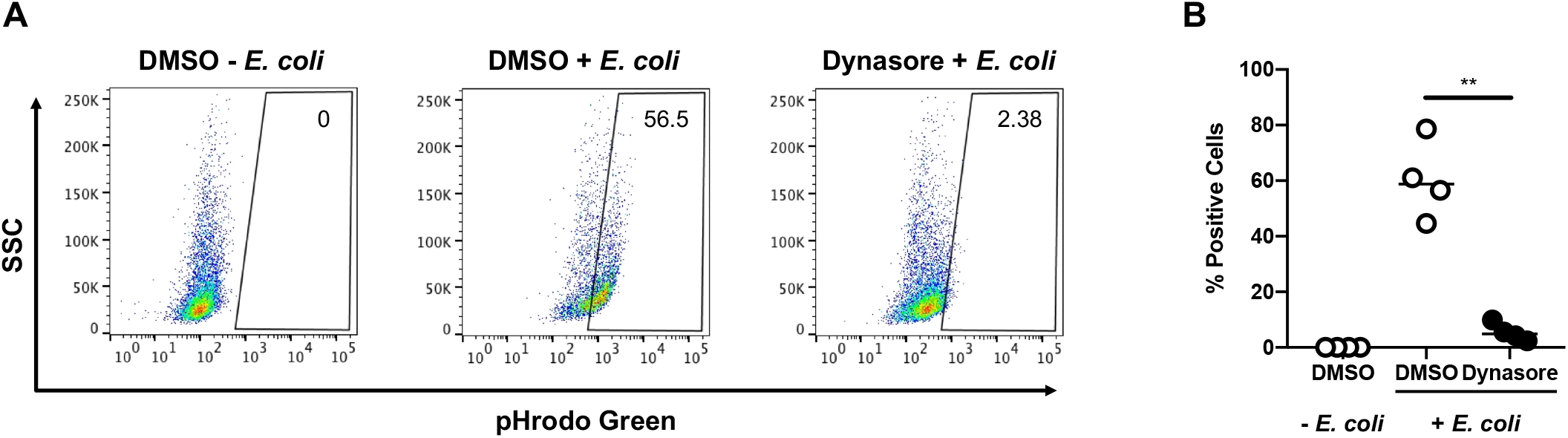
Dynasore inhibits endocytosis/phagocytosis of labeled *E. coli* particles by MDMs. (A-B) MDMs (5×10^5^/well) were incubated with DMSO or Dynasore (80 μM) for 15 minutes at 37°C. The cells were washed with PBS and incubated with pHrodo Green *E. coli* (1 mg/ml) for 2 hours at 37°C. Endocytosis/phagocytosis was measured by flow cytometry. Shown are data from one representative donor (A) and composite data from four donors (B). *, p < 0.05; **, p < 0.01; ***, p < 0.001; p < 0.0001.

**Supplemental Figure 5.**
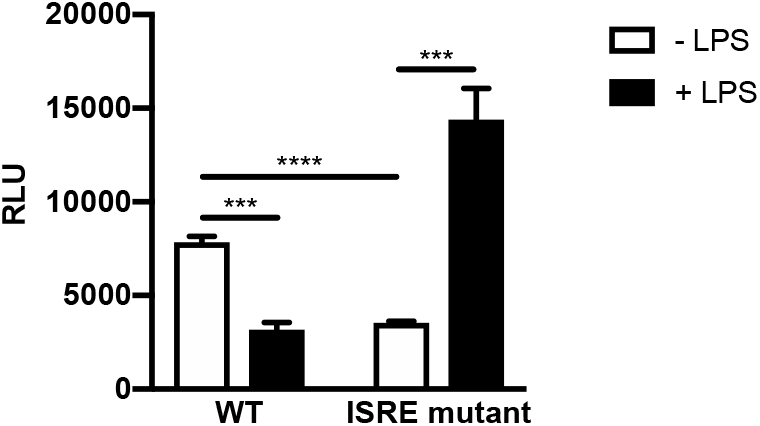
TLR4-mediated repression of HIV-1 requires the ISRE binding site located in the GLS downstream of the 5′ LTR. (A) HEK293-TLR4^CFP^/MD-2/CD14 cells were transfected with HIV-1 LTR/GLS-luciferase reporter constructs with an intact ISRE or mutated ISRE binding site. Following transfection, cells were treated with LPS (100 ng/ml) for 18 hours and then harvested and assayed for luciferase activity. Data are the mean (± SD) of three independent experiments, each performed in triplicate. *, p < 0.05; **, p < 0.01; ***, p < 0.001; p < 0.0001.

